# Aging Boosts Antiviral CD8^+^T Cell Memory Through Improved Engagement Of Diversified Recall Response Determinants

**DOI:** 10.1101/450072

**Authors:** Bennett Davenport, Jens Eberlein, Tom T. Nguyen, Francisco Victorino, Kevin Jhun, Haedar Abuirqeba, Verena van der Heide, Peter Heeger, Dirk Homann

## Abstract

The determinants of protective CD8^+^ memory T cell (CD8^+^T_M_) immunity remain incompletely defined and may in fact constitute an evolving agency as aging CD8^+^T_M_ progressively acquire enhanced rather than impaired recall capacities. Here, we show that old as compared to young antiviral CD8^+^T_M_ more effectively harness disparate molecular processes (cytokine signaling, trafficking, effector functions, and co-stimulation/inhibition) that in concert confer greater secondary reactivity. The relative reliance on these pathways is contingent on the nature of the secondary challenge (greater for chronic than acute viral infections) and over time, aging CD8^+^T_M_ re-establish a dependence on the same accessory signals required for effective priming of naïve CD8^+^T cells in the first place. Thus, our findings are consistent with the recently proposed “rebound model” that stipulates a gradual alignment of naïve and CD8^+^T_M_ properties, and identify a diversified collection of potential targets that may be exploited for the therapeutic modulation of CD8^+^T_M_ immunity.

## INTRODUCTION

What does it take for pathogen-specific CD8^+^ memory T cells (CD8^+^T_M_) to mount an efficient and protective recall response? In most general terms, the efficacy of a secondary (II^o^) CD8^+^ effector T cell (CD8^+^T_E_) response is contingent on the numbers of available CD8^+^T_M_, their differentiation status and anatomical distribution, the contribution of other immune cell populations (*e.g.,* CD4^+^T cells, B cells, innate immune cells), and the precise conditions of pathogen re-encounter, *i.e.* the nature of the pathogen as well as the route and dosage of infection. Thus, the specific constraints of experimental or naturally occurring pathogen exposure will dictate relevant outcomes that are predictable only in as much as the relative contribution of individual biological parameters are sufficiently understood, a task much complicated by the considerable combinatorial possibilities that ultimately shape the balance of pathogen replication and control, pathogen-induced damage, immunopathology, tissue protection and repair. Simply put, CD8^+^T_M_-mediated immune protection is eminently context-dependent.

The difficulties associated with attempts to define more generally applicable rules for the phenomenon of protective CD8^+^T_M_ immunity are perhaps best illustrated by the “effector/central memory T cell” paradigm (T_EM_ and T_CM_, respectively) that constitutes one of the most widely employed and consequential distinctions in the field of memory T cell research [1]. The analytical and physical separation according to CD62L (and CCR7) expression status has spawned an extraordinary amount of work that has assigned numerous distinctive, and at times seemingly contradictory, properties to CD62L^lo^ CD8^+^T_EM_ and CD62L^hi^ CD8^+^T_CM_ subsets [2-4]. The CD8^+^T_M_ populations thus defined, however, are very much a moving target. For example, CD62L expression by peripheral CD8^+^T_M_ generated in response to an acute pathogen challenge is progressively enhanced as a function of original priming conditions and infection history; upon entry into certain lymphoid or nonlymphoid tissues, CD8^+^T_M_-expressed CD62L is reduced; and CD8^+^T_EM_ and T_CM_ subsets themselves are subject to a gradual adaptation that introduces an array of molecular, phenotypic and functional changes including, importantly, an increase of their respective recall capacities [2, 5-10]. Most recently, D. Busch’s group used an elegant serial adoptive transfer system in which single I^o^, II^o^ or III^o^ *L. monocytogenes-* (LM-) specific CD8^+^T_CM_ (*i.e.*, CD8^+^T_CM_ established after a I^o^, II^o^ or III^o^ LM challenge) gave rise to recall responses of comparable size, phenotypic and functional diversity, and protective capacity [11, 12]. Since single CD8^+^T_EM_ failed to mount a similar response, these studies provide definitive proof that the CD62L^hi^ CD8^+^T_CM_ subset harbors greater recall potential [11, 12] yet CD62L itself is apparently dispensable for an effective LM-specific recall response [13]. In some other model systems, enhanced protection was even afforded by CD8^+^T_EM_, their limited proliferative potential notwithstanding [2-4]. It is therefore imperative to define, beyond the T_EM_/T_CM_ paradigm, which exact mechanisms contribute to the regulation of effective CD8^+^T_M_ recall activity under varied experimental conditions, and to what extent specific molecular pathways may become a dominant force in a given model system. A synthesis of such efforts may then provide a foundation for the formulation of more general rules of CD8^+^T_M_ engagement.

In the present work, we took advantage of our observation that aging CD8^+^T_M_ specific for lymphocytic choriomeningitis virus (LCMV) gradually acquire unique molecular, phenotypic and functional signatures that are associated with a capacity for more vigorous II^o^ CD8^+^T_E_ responses and improved immune protection [9]. We have further organized these dynamic changes in the “rebound model” of extended CD8^+^T_M_ maturation according to which pertinent properties of aging CD8^+^T_M_ are progressively aligned, perhaps surprisingly, with those of naïve CD8^+^T_N_ populations [9, 10]. Here, by focusing on a diverse set of co-stimulatory and inhibitory, cytokine, chemokine and homing receptors/ligands differentially expressed by old and young CD8^+^T_M_ as well as their distinct effector function profiles [9], we identified a broad array of mechanisms that “tune” CD8^+^T_M_ recall reactivity to an acute and/or chronic viral re-challenge, and that specifically support the greater II^o^ CD8^+^T_E_ expansions of aged CD8^+^T_M_ populations. In particular, we propose that aging CD8^+^T_M_ re-acquire a dependence on multiple accessory pathways for optimization of their II^o^ CD8^+^T_E_ reactivity that were essential for the effective and efficient priming of naïve CD8^+^T_N_ in the first place.

## RESULTS

### Interrogating CD8^+^T_M_ recall responses: the mixed adoptive transfer/re-challenge (AT/RC) system

To identify the mechanisms regulating the differential recall reactivity of young and old antiviral CD8^+^T_M_, we employed a mixed “adoptive transfer/re-challenge” (AT/RC) system described in ref.[9]. In brief, cohorts of young adult mice congenic at the CD45 or CD90 locus were challenged with LCMV (2×10^5^ pfu LCMV Armstrong [Arm] i.p.) and allowed to establish LCMV-specific CD8^+^T cell memory. By performing viral infections in a staggered fashion, we generated groups of young (∼2 months after challenge) and aged (>15 months after infection) LCMV-immune mice that served as donors for a concurrent interrogation of young and old CD8^+^T cell memory. To this end, CD8^+^T_M_ populations were enriched from the congenic donors, combined at a ratio of 1:1 at the level of CD8^+^T_M_ specific for the immunodominant LCMV nucleoprotein (NP) determinant NP_396-404_ (D^b^NP_396_^+^CD8^+^T_M_), and injected into congenic recipients that were subsequently inoculated with LCMV; the respective expansions of young *vs.* old D^b^NP_396_^+^CD8^+^T_M_-derived II^o^ CD8^+^T_E_ populations were then quantified eight days later (***Fig.1A***).

**Figure 1.**
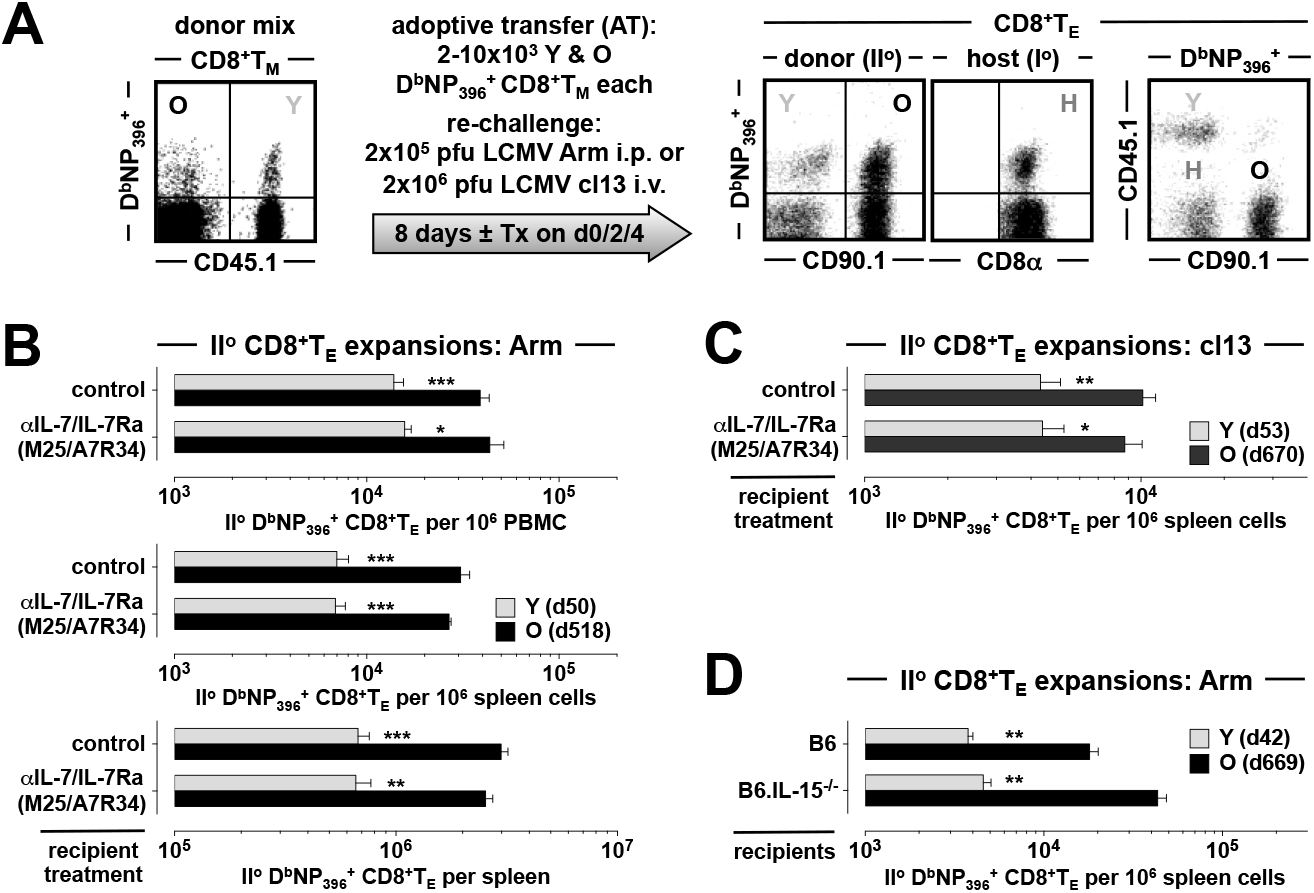
No role for IL-7, TSLP or IL-15 in the differential regulation of young and old II^o^ CD8^+^T_E_ expansions. **A.,** basic design of mixed AT/RC experiments. CD8^+^T cells from congenic young and old LCMV-immune donors were enriched, combined 1:1 at the level of D^b^NP_396_^+^CD8^+^T_M_, and transferred i.v. into recipients that were subsequently challenged using “acute” (LCMV Arm) or “chronic” (LCMV cl13) infection protocols; proliferative expansions of II^o^ D^b^NP_396_^+^CD8^+^T_E_ were quantified 8 days later. Note that the constellation of congenic markers permits the distinction of young and old II^o^ CD8^+^T_E_ as well as I^o^ CD8^+^T_E_ generated by the host. Unless noted otherwise, treatment with blocking antibodies was performed ∼2h before AT and on d2 and d4 after virus inoculation; in other cases, B6 *vs*. immunodeficient recipients were used (not shown). **B.,** quantification of II^o^ CD8^+^T_E_ expansions under conditions of control (PBS) or combined αIL-7/αIL-7Ra treatment and LCMV Arm challenge; the age of donor CD8^+^T_M_ is indicated in the legend (young: d50, old: d518) **C.,** similar experiments as in panel B but conducted with LCMV cl13 and control treatment with rat IgG (donor age indicated in legend). **D.,** mixed AT/RC experiments performed with B6 *vs*. B6.IL-15^-/-^ recipients (AT of 2×10^3^ [panel B & D] or 10×10^3^ [panel C] young and old D^b^NP_396_^+^CD8^+^T_M_ each; n≥3 mice/group; asterisks indicate significant differences comparing young and old II^o^ D^b^NP_396_^+^CD8^+^T_E_ populations using Student’s t-test.

As detailed in ref.[9], the mixed AT/RC model offers several practical advantages that facilitate the elucidation of molecular mechanisms in control of differential CD8^+^T_M_ recall capacities. 1., young and old II^o^ CD8^+^T_E_ responses develop in the same host and are therefore subject to the same general perturbations provoked by various experimental interventions. 2., the magnitude of recall responses elaborated by transferred CD8^+^T_M_ populations is primarily shaped by a complex of T cell-intrinsic properties and, importantly, is largely independent of host age. 3., the relative extent of II^o^ CD8^+^T_E_ expansions (but not the functional profiles of II^o^ CD8^+^T_E_) can serve as a correlate for immune protection. 4., the AT of low CD8^+^T_M_ numbers permits their maximal *in vivo* activation without preventing the generation of concurrent I^o^ CD8^+^T_E_ responses; accordingly, the system can monitor the relatively independent evolution of three CD8^+^T_E_ populations targeting the same viral epitope (I^o^, young II^o^ and old II^o^ D^b^NP_396_^+^CD8^+^T_E_; ***Fig.1A***). 5., the use of two different re-challenge protocols may differentiate between basic determinants required for CD8^+^T_M_ recall responses in the wake of an “acute” LCMV Arm infection (AT/RC Arm) and a more complex constellation of mechanisms supporting the effective coordination II^o^ CD8^+^T_E_ expansions after a “chronic” LCMV clone 13 infection (AT/RC cl13) (***Fig.1A***). Altogether, we deployed the mixed AT/RC approach to ascertain the contribution of particular molecular pathways to the divergent II^o^ expansion of young and old CD8^+^T_M_ by treatment of recipients with blocking antibodies or use of immunodeficient hosts (***Fig.1A***); while the systemic nature of these interventions cannot discern between direct and indirect effects exerted on CD8^+^T cell populations, the broad utility and practical relevance of our approach lies in the relative ease with which CD8^+^T_E_ cell responses can be reliably manipulated. Lastly, for facilitated manipulation of CD8^+^T_M_ we employed the “p14 chimera” model in which purified naïve and congenic p14 T_N_ (TCRtg CD8^+^T cells specific for LCMV glycoprotein GP_33-41_) are transferred into B6 recipients that are subsequently challenged with LCMV Arm to generate young and old p14 T_M_ [9] (since p14 T_M_ are a clonotypic population, p14 chimeras also effectively control for TCR affinity/avidity as a potentially confounding variable).

### No role for IL-7 and IL-15 in the differential regulation of young and old II^o^ CD8^+^T_E_ expansions

The cytokines IL-7 and IL-15 are essential for the preservation of CD8^+^T cell memory as they support homeostatic proliferation and survival of CD8^+^T_M_ [14]. Accordingly, the pronounced upregulation of IL-7 and IL-15 receptor components (CD127/IL-7Ra and CD122/IL-2Rb) by aging CD8^+^T_M_ suggested their increasing responsiveness to IL-7 and IL-15 (ref.[9] and ***Fig.S1*** which stratifies an enrichment of JAK-STAT pathway genes in old p14 T_M_). Although this is indeed the case (determined at the level of cytokine-induced STAT5 phosphorylation), homeostatic proliferation rates of antiviral CD8^+^T_M_ remained surprisingly unaffected by age [10]. Nevertheless, enhanced CD8^+^T_M_ expression of CD127 and CD122 could still contribute to improved recall responses since both IL-7 and IL-15 may act as “adjuvants” to boost CD8^+^T_E_ immunity [15, 16], albeit in a potentially STAT5-independent manner for II^o^ CD8^+^T_E_ responses [17]. We therefore employed the mixed AT/RC system (***Fig.1A***) to evaluate the impact of combined IL-7/IL-7Ra blockade on the II^o^ reactivity of young and old CD8^+^T_M_. As shown in ***Fig.1B***, both differential and overall II^o^ CD8^+^T_E_ expansions after an “acute” LCMV Arm challenge were impervious to IL-7/IL-7Ra blockade; the data also illustrate that an analysis of different tissues (blood or spleen) and the use of different denominators (II^o^ CD8^+^T_E_ per 10^6^ cells or total spleen cells) provides essentially similar results (***Fig.1B***). Similarly, IL-7/IL-7Ra blockade remained without consequences in additional mixed AT/RC experiments using the “chronic” LCMV cl13 model (***Fig.1C***). Our results further exclude a relevant contribution of thymic stromal lymphopoietin (TSLP) to II^o^ CD8^+^T_E_ expansions since the TSLP receptor (TSLPR), downregulated by aging CD8^+^T_M_ (***Fig.S1*** and ref.[9]), associates with CD127 for effective signal transduction [18], and the CD127-specific A7R34 antibody used in our experiments also inhibits TSLP action [19].

We also ascertained a potential role for IL-15 in our model system by conducting mixed AT/RC experiments with IL-15^-/-^ recipients. Lack of IL-15, however, did not compromise the greater II^o^ reactivity of old CD8^+^T_M_ (***Fig.1D***); in fact, recall responses of aged CD8^+^T_M_ were somewhat increased in IL-15^-/-^ as compared to B6 control mice (2.4-fold, p=0.01; ***Fig.1D***), perhaps as a result of lymphopenia-enhanced, IL-15-independent expansions in the context of an active inflammation. We conclude that elevated CD127 and CD122 expression by aging CD8^+^T_M_ does not improve their recall responses.

### Divergent requirements of IL-4, IL-6 and TGFβ for enhanced II^o^ reactivity of aged CD8^+^T_M_

The dynamic regulation of CD127 and CD122 expression discussed above constitutes a common theme shared by multiple other CD8^+^T_M_-expressed cytokine receptors. With the notable exception of TSLPR, these receptors are all subject to increasing expression by aging CD8^+^T_M_ with overall gains varying from the modest (CD126/IL-6Ra, CD130/IL-6ST, IL-21R, IFNAR1) to the more pronounced (CD124/IL-4Ra, TGFβRII, CD119/IFNγR1) (***Fig.S1*** and ref.[9]). Corresponding temporal analyses extended here to blood-borne CD8^+^T_M_ populations with different LCMV specificities further support the conclusion that the prolonged phenotypic CD8^+^T_M_ maturation is indeed a generalized and systemic phenomenon (***Figs.2A/B & S2***). The kinetics of CD124, CD126 and TGFβRII expression are of particular interest since the signaling pathways downstream of these receptors emerged as distinctive traits in our earlier Ingenuity Pathway Analyses of aging CD8^+^T_M_ [9], and both IL-6 and TGFβ have been suggested to exert crucial roles in the natural history of chronic LCMV infection [20, 21]. To further assess the relation between cytokine receptor expression levels and signal transduction capacity, we briefly exposed young and old p14 T_M_ *in vitro* to IL-4 or IL-6 and quantified phosphorylation of STAT6 and STAT3, respectively. Here, aged p14 T_M_ indeed responded with greater STAT phosphorylation, and the re-expression of CD124 by old p14 T_M_ at levels otherwise found only on naïve CD8^+^T cells correlated with equal IL-4 reactivity of these populations (***Fig.2B***). The generally lower CD126 (and CD130 [9]) expression by CD8^+^T_M_, which required overall higher cytokine concentrations for effective STAT phosphorylation as compared to the IL-4 experiments, nevertheless conferred an age-dependent differential induction of pSTAT3; at the same time, IL-10-induced STAT3 phosphorylation demonstrated no differences (***Fig.2B***) in agreement with the stable low-level IL-10 receptor expression among aging CD8^+^T_M_ [9].

**Figure 2.**
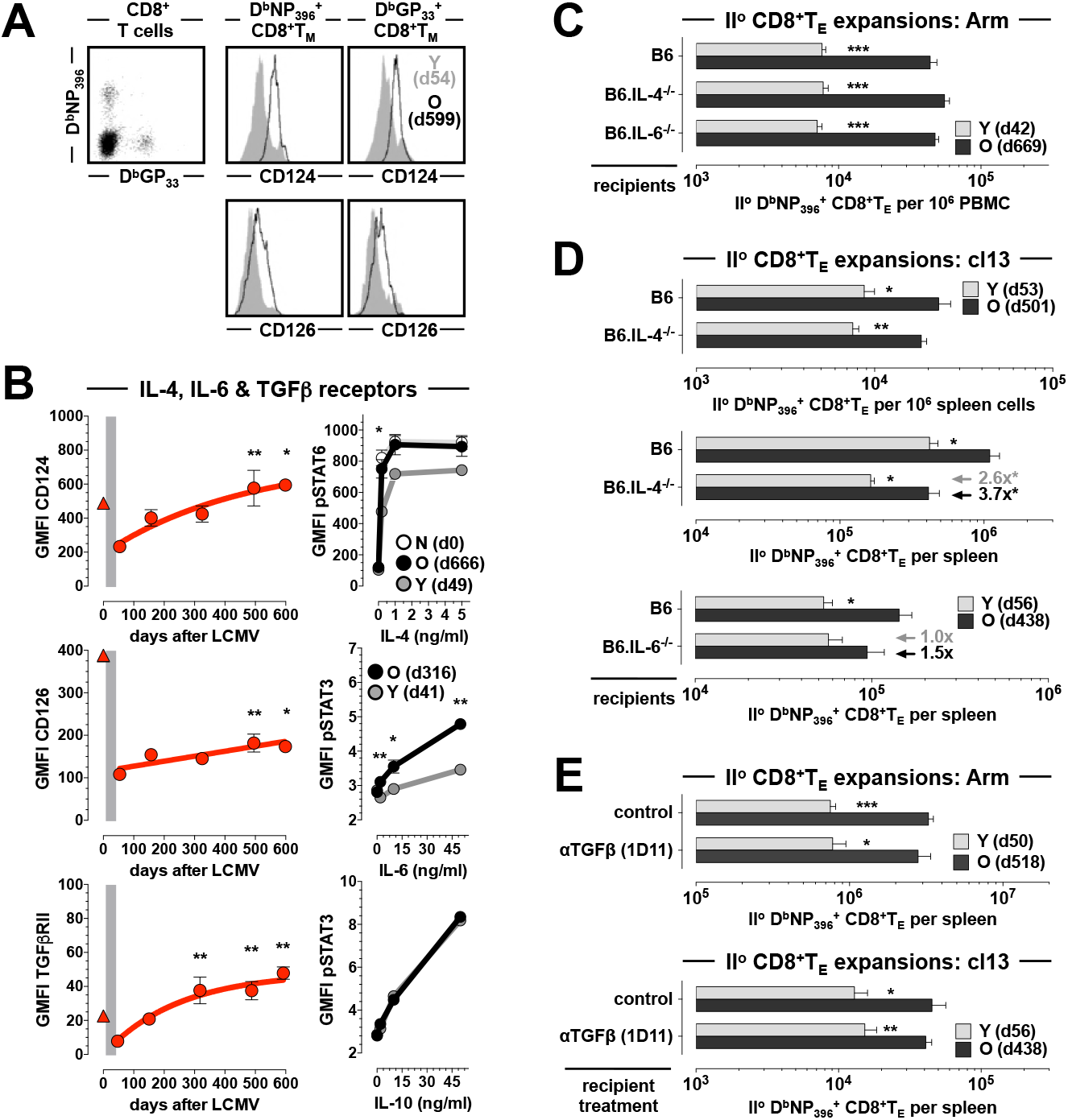
Divergent requirements of IL-4, IL-6 and TGFb for enhanced II^o^ reactivity of aged CD8^+^T_M_. **A.,** cytokine receptor expression levels by blood-borne D^b^NP_396_^+^ and D^b^GP_33_^+^CD8^+^T_M_ (left plot) were quantified in contemporaneous analyses of aging LCMV-immune mice by determining their respective GMFI values (geometric mean of fluorescent intensity); the overlaid histograms depict representative CD124 and CD126 expression by young (gray) and aged (black tracing) D^b^NP_396_^+^ (middle) and D^b^GP_33_^+^ (right) CD8^+^T_M_. **B.,** left plots: temporal regulation of CD124, CD126 and TGFβRII expression by aging D^b^NP_396_^+^CD8^+^T_M_ (triangle symbol: naïve CD44^lo^CD8^+^T_N_; the gray bar demarcates the period from peak I^o^ CD8^+^T_E_ expansion [d8] to initial establishment of CD8^+^T cell memory [d42], and asterisks indicate statistical significance comparing young and older D^b^NP_396_^+^CD8^+^T_M_ using one-way ANOVA with Dunnett’s multiple comparisons test). Right plots: STAT phosphorylation by young (gray) and old (black) p14 T_M_ was assessed directly *ex vivo* and after 15min *in vitro* culture in the presence of graded dosages of recombinant IL-4 (top), IL-6 (middle) or IL-10 (bottom); the top panel also includes an analysis of naïve p14 T_N_ (white). **C.,** II^o^ CD8^+^T_E_ expansions in B6, B6.IL-4^-/-^ and B6.IL-6^-/-^ mice after mixed AT/RC Arm. **D.,** similar experiments as in panel B but performed with LCMV cl13. **E.,** II^o^ CD8^+^T_E_ expansions under conditions of TGFβ blockade. The gray and black arrows/values in panel C indicate the extent of significantly reduced (asterisks) II^o^ CD8^+^T_E_ expansions comparing young II^o^ CD8^+^T_E_ in B6 and B6.IL-4^-/-^ mice (gray), as well as old II^o^ CD8^+^T_E_ in B6 and B6.IL4^-/-^mice (black) (n≥3 mice/group; AT of 2×10^3^ [panel C & E top], 10×10^3^ [panel D top/middle] or 5×10^3^ [panel D bottom & E bottom] young and old D^b^NP_396_^+^CD8^+^T_M_ each).

Despite the heightened reactivity of old CD8^+^T_M_ to IL-4, initial experiments performed with the mixed AT/RC Arm approach and B6 *vs.* IL-4^-/-^ recipients did not reveal a role for IL-4 in the regulation of II^o^ CD8^+^T_E_ expansions (***Fig.2C***). In contrast, LCMV cl13 infection of IL-4^-/-^ recipients resulted in an overall decrease of specific CD8^+^T_E_ immunity, including a 4.0-fold reduction of the splenic I^o^ CD8^+^T_E_ response (p=0.0056). Importantly, the relative reduction of old II^o^ CD8^+^T_E_ expansions was more pronounced (3.7-fold, comparing B6 and IL-4^-/-^ recipients) than that of respective young II^o^ CD8^+^T_E_ populations (2.6-fold) (***Fig.2D***, note arrows, values [3.7x *vs.* 2.6x], and significance [asterisks]). Collectively, these findings add to an emerging consensus about the importance of IL-4 for the generation of effective antiviral CD8^+^T cell immunity [22, 23] and demonstrate a specific requirement for IL-4 to support the greater II^o^ reactivity of aged CD8^+^T_M_; the direct correlation between CD124 expression levels of CD8^+^T_M_ and their recall potential as well as the comparable reduction of I^o^ and old II^o^ CD8^+^T_E_ expansions in IL-4^-/-^ mice are further consistent with predictions of the “rebound model” that a progressive alignment of CD8^+^T_N_ and aging CD8^+^T_M_ properties may translate into a reliance on similar co-stimulatory requirements [9].

IL-6 is among the most prominent cytokines induced after an LCMV infection [24] but despite the enhanced responsiveness of aged CD8^+^T_M_ to IL-6 stimulation (***Fig.2B***), the differential II^o^ responses of transferred young and old CD8^+^T_M_ were not compromised by an LCMV Arm challenge of IL-6^-/-^ recipients (***Fig.2C***). Using the LCMV cl13 infection protocol, IL-6-deficiency imparted a very modest 1.5-fold reduction of aged but not young II^o^ CD8^+^T_E_ expansions that also mirrored a 1.4-fold decrease of the I^o^ response; neither finding, however, proved significant (***Fig.2D*** and not shown) suggesting an overall more limited contribution of IL-6 to differential young and old CD8^+^T_M_ recall immunity. As to the potential function of TGFβ and related cytokines in the context of CD8^+^T_M_ aging, we earlier noted a series of marked transcriptional adaptations (increasing mRNA abundance for activin and BMP receptors as well as *Smad1*) and further identified a pronounced increase of TGFβRII protein (but not mRNA) expression by aging CD8^+^T_M_ (***Figs.2B, S2*** and ref.[9]). The functional relevance of TGFβRII-mediated signaling for LCMV-specific CD8^+^T cells has been illustrated in a persistent infection model where suppression of TGFβRII activity specifically in T cells improved virus control [20]. Thus, increased TGFβRII expression by aging CD8^+^T_M_ could conceivably operate to constrain exuberant II^o^ CD8^+^T_E_ responses. More recent work, however, could not demonstrate a therapeutic effect of TGFβ blockade in chronic LCMV infection [25, 26], and in agreement with those studies we did not observe an unleashing of old II^o^ CD8^+^T_E_ immunity in our mixed AT/RC system following TGFβ blockade, nor could we discern any impact on the CD8^+^T_M_ recall responses at large in either acute or chronic infection models (***Fig.2E***).

### Contributions of IFNγ, IFNγ receptor and FasL to the differential regulation of CD8^+^T_M_ recall responses

In addition to multiple phenotypic alterations, aging of CD8^+^T_M_ also introduces a number of changes among their functional properties that collectively foster a more diversified spectrum of effector activities [9]. Notably, old CD8^+^T_M_ produce more IFNγ on a per cell basis, and a greater fraction of aged CD8^+^T_M_ can be induced to express Fas ligand (FasL) [9]. Together with IL-2, the production capacity of which modestly increases with age [9, 27], IFNγ and FasL also share the distinction as the only CD8^+^T_M_ effector molecules whose cognate receptors (CD122, CD119 and CD95/Fas, respectively) are concurrently upregulated by aging CD8^+^T_M_ (***Fig.S1*** and refs.[9, 10]). This can have direct implications for the autocrine regulation of CD8^+^T_M_ immunity in the context of recall responses as documented for IL-2 [28], and similar considerations may also apply to IFNγ given that its direct action on CD8^+^T cells is required for optimal I^o^ CD8^+^T_E_ expansions and CD8^+^T_M_ development [29]. If CD8^+^T_M_-intrinsic interactions between FasL and Fas shape II^o^ CD8^+^T_E_ immunity, however, remains elusive.

To correlate the differential CD119 expression by young and old CD8^+^T_M_, confirmed and extended here to different LCMV-specific CD8^+^T_M_ populations in peripheral blood (***Figs.3A & S2***), with a direct responsiveness to IFNγ action, we determined the extent of STAT1 phosphorylation in young and old p14 T_M_. Interestingly, aged p14 T_M_ featured a slight yet significant elevation of constitutive STAT1 phosphorylation, a difference that was further amplified by *in vitro* exposure to IFNγ (***Fig.3A***). Thus, taking into account differential CD119 expression levels, responsiveness to IFNγ, and IFNγ production capacities of young and old CD8^+^T_M_ [9], we conducted a first set of mixed AT/RC experiments with IFNγ^-/-^ recipients. In this system, IFNγ production is restricted to the transferred CD8^+^T_M_ populations but both host cells and donor CD8^+^T_M_ can readily respond to IFNγ. Comparing CD8^+^T_M_ recall responses in LCMV Arm-infected B6 *vs.* IFNγ^-/-^ recipients, we found that absence of host IFNγ modestly compromised the II^o^ expansions of both young and old CD8^+^T_M_, though unexpectedly the relative decrease was more pronounced for the former rather than the latter population (***Fig.3B***). We therefore extended our experiments to assess the contribution of IFNγ at large by use of a neutralizing antibody. Here, complete IFNγ blockade further reduced II^o^ CD8^+^T_E_ responses and in particular impaired the II^o^ response of aged CD8^+^T_E_ (***Fig.3B***; compare the increase of relative reductions among young II^o^ CD8^+^T_E_ expansions [blood: from 2.6x in IFNγ^-/-^ hosts to 2.9x after IFNγ blockade; a 1.1x increase] to those of aged II^o^ CD8^+^T_E_ populations [blood: from 1.5x in IFNγ^-/-^ hosts to 2.4x after IFNγ blockade; a 1.6x increase]). Together, our findings demonstrate a moderate role for IFNγ in the regulation of CD8^+^T_M_ recall responses to an acute LCMV challenge that differs according to the cellular source of IFNγ: while the II^o^ expansion of young CD8^+^T_M_, despite reduced CD119 expression and signaling, is more reliant on IFNγ production by other cells, aged CD8^+^T_M_ populations, on account of enhanced IFNγ production capacity [9], can better promote their own II^o^ reactivity. This notion is further reinforced by mixed AT/RC experiments using LCMV cl13 infection under conditions of IFNγ blockade. As shown in ***Fig.3C***, neutralization of IFNγ profoundly depressed II^o^ CD8^+^T_E_ immunity and largely extinguished any differences between young and old II^o^ CD8^+^T_E_ expansions (note the comparable population sizes in blood [top] and at the level of total II^o^ CD8^+^T_E_ per spleen [bottom] in ***Fig.3C***).

**Figure 3.**
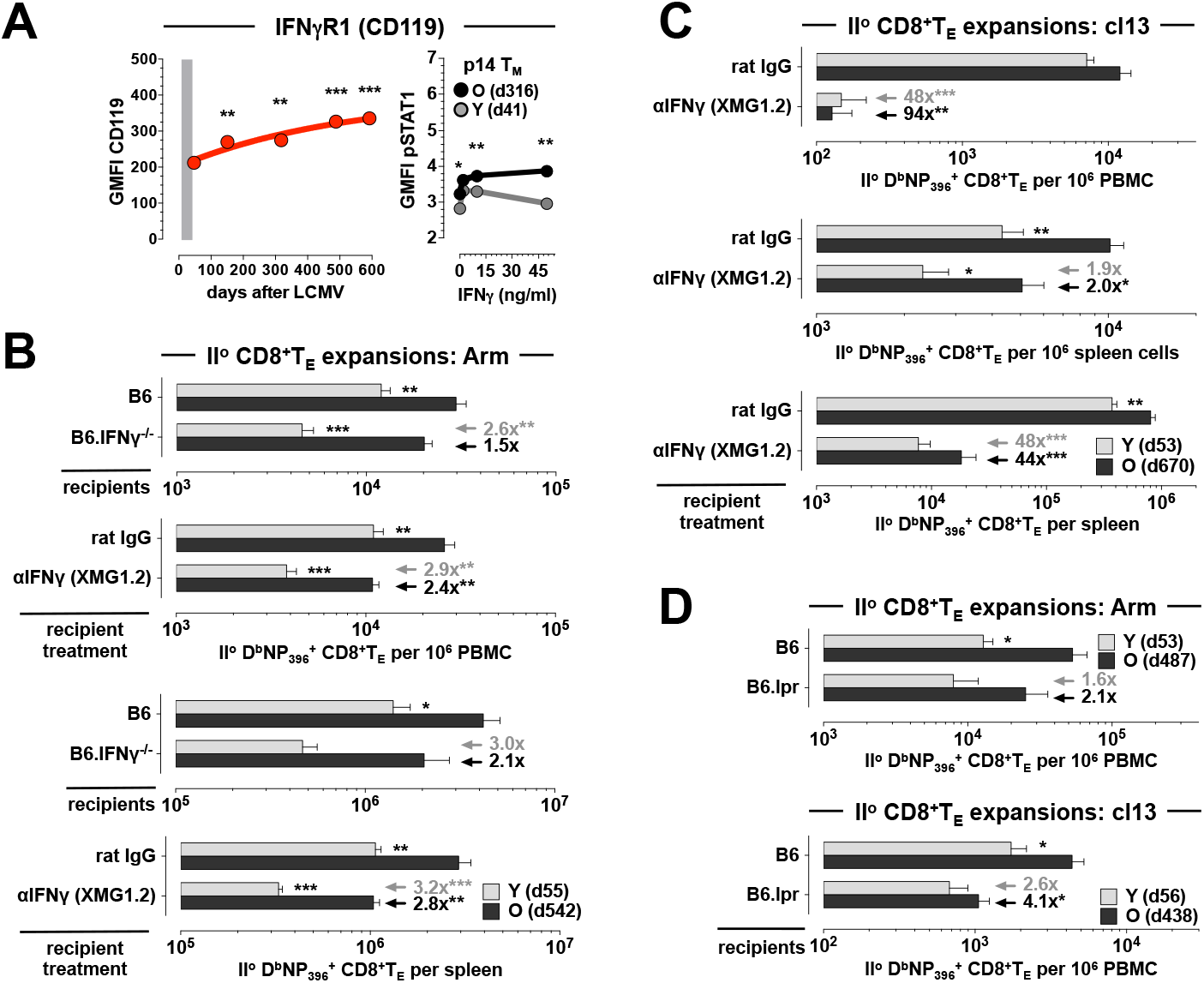
Role of IFNγ, IFNγ receptor and FasL in the regulation of young and old CD8^+^T_M_ recall activity. **A.,** left: expression of CD119 by aging D^b^NP_396_^+^CD8^+^T_M_ in the PBMC compartment. Right: STAT1 phosphorylation by young and aged p14 T_M_ was determined *ex vivo* and after 15min *in vitro* exposure to recombinant IFNγ; note the slightly enhanced *ex vivo* pSTAT1 levels in old *vs*. young p14 T_M_. **B.,** mixed AT/RC Arm experiments were performed with B6 and B6.IFNγ^-/-^ recipients as well as under conditions of control (rat IgG) or aIFNg treatment. **C.,** similar IFNγ blocking experiments as in panel B but conducted with the chronic LCMV cl13 model. **D.,** II^o^ CD8^+^T_E_ expansions after AT/RC Arm (top) or AT/RC cl13 (bottom) using B6 *vs*. B6.lpr recipients. Arrows/values in panel B-D indicate the respective extent and significance (asterisks) by which antibody treatment or immunodeficiency reduced II^o^ expansions of young (gray) or old (black) II^o^ CD8^+^T_E_ populations (n≥3 mice/group or time point; AT of 3×10^3^ [panel B], 10×10^3^ [panel C], 2×10^3^ [panel D top] or 5×10^3^ [panel D bottom] young and old D^b^NP_396_^+^CD8^+^T_M_ each).

In contrast to IFNγ, the role of FasL:Fas interactions in the LCMV model appears more limited - both FasL- and Fas-mutant mice (FasL^gld^ and Fas^lpr^ strains, respectively) control an acute LCMV infection [30] - yet a non-redundant role for Fas in virus clearance or CD8^+^T_M_ generation could be readily demonstrated in mice with compound immunodeficiencies [31-33]. To evaluate the contribution of the FasL:Fas pathway in our model system, we conducted mixed AT/RC experiments with B6 *vs.* Fas^lpr^ (“B6.lpr”) recipients and observed a preferential reduction of aged II^o^ CD8^+^T_E_ expansions in the B6.lpr hosts that was especially pronounced following chronic LCMV cl13 infection (***Fig.3D***). Although we can conclude that the enhanced II^o^ reactivity of old CD8^+^T_M_ is in part controlled by their broader FasL induction, the precise mechanisms operative in this context remain to be elucidated and may involve accelerated virus clearance [9] through FasL-dependent cytolysis, nonapoptotic FasL:Fas interactions between CD8^+^T_M_ and T_N_ that facilitate concurrent I^o^ CD8^+^T_E_ differentiation [34], or perhaps the autocrine binding of secreted FasL that, akin to a mechanism proposed for tumor cells [35], may shield II^o^ CD8^+^T_E_ from FasL-mediated fratricide.

### LFA-1 and CXCR3 blockade preferentially curtail II^o^ expansions of aged CD8^+^T_M_

Among the array of phenotypic changes accrued during CD8^+^T_M_ aging we previously noted several cell surface receptors involved in the regulation of CD8^+^T cell traffic and migration [9] and demonstrated the importance of their age-associated expression differences in the context of immune homeostasis [10]. Now, using an unbiased approach based on time series gene set enrichment analyses (GSEA) of aging p14 T_M_ populations [10], the potential importance of differential “homing receptor” expression is further supported by our identification of the “cell adhesion molecules” module as the top KEGG pathway negatively enriched in old p14 T_M_ (normalized enrichment score: −1.82; p=0.0078; ***Fig.4A***). For 29/38 genes within this module, we also performed temporal protein expression analyses and demonstrated a significant up- or downregulation by aging CD8^+^T_M_ for half of these gene products (15/29; ***Fig.4A*** and ref.[9]). Here, the expression pattern of CD11a/integrin α_L_ caught our attention for several reasons: elevated CD11a expression, similar to CD44, has long been used as a surrogate marker for “antigen-experienced” CD8^+^T cells [36]. In combination with CD18/integrin β_2_, CD11a forms the heterodimeric LFA-1 complex that constitutes, together with its endothelial cell-expressed ligands CD54/ICAM1 and CD102/ICAM2, one of the major pathways for leukocyte adhesion. In contrast to CD44, however, CD11a mRNA and protein expression by aging CD8^+^T_M_ are subject to a slight yet significant decline (***Figs.4A/B, S2*** and ref.[9]). In fact, other components of the LFA-1 pathway exhibited very similar patterns with a progressive decrease of CD8^+^T_M_-expressed CD18, CD102 and in particular CD54 mRNA and/or protein; another LFA-1 ligand, CD50/ICAM5, is not expressed by murine CD8^+^T_E/M_ (***Fig.4A/B, S2*** and ref.[9]).

**Figure 4.**
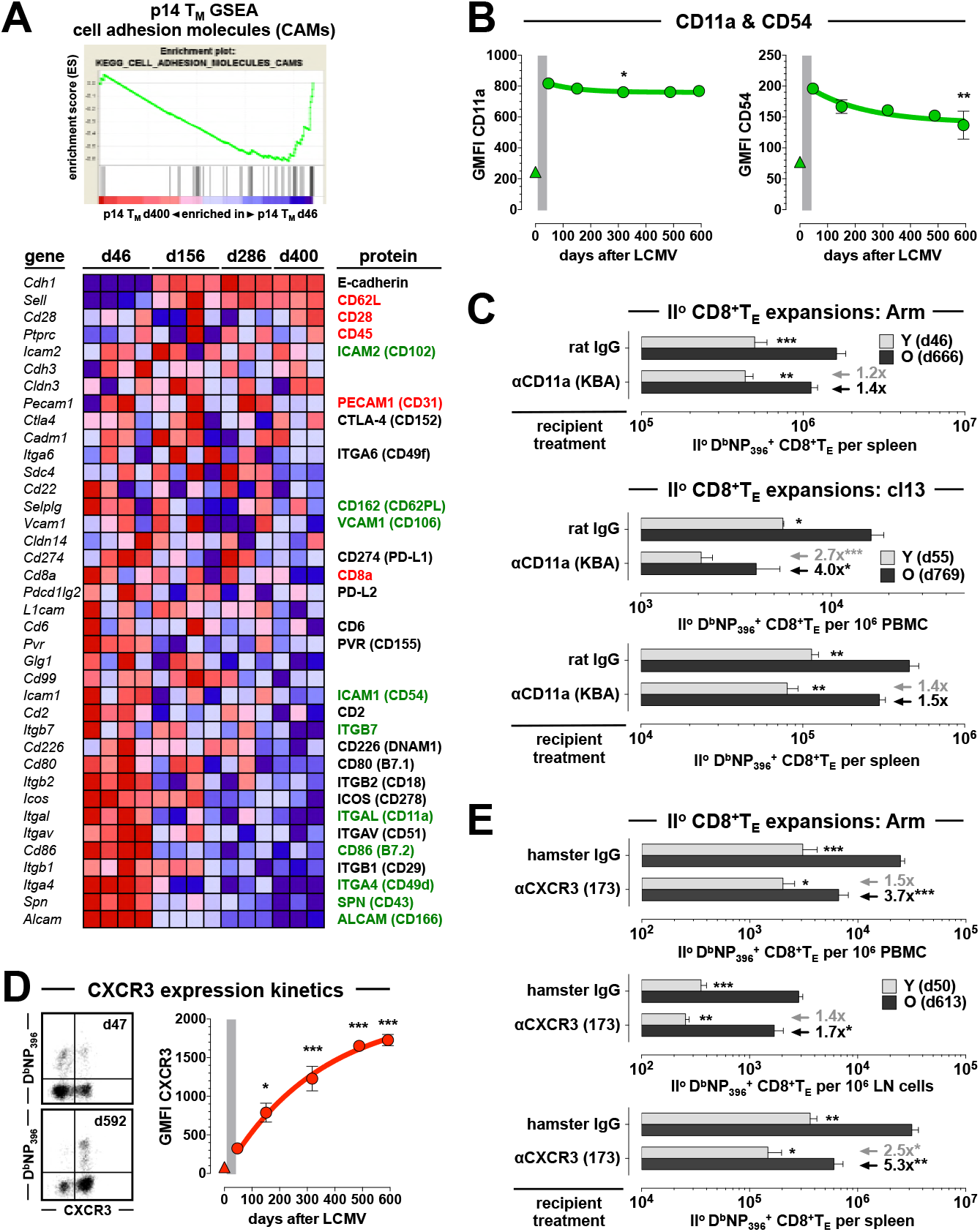
CD11a and CXCR3 blockade preferentially restrict II^o^ expansions of aged antiviral CD8^+^T_M_. **A.,** time series GSEA were conducted for *ex vivo* purified aging p14 T_M_ (d46, d156, d286 and d400) as detailed in refs. [9, 10]. Top: old p14 T_M_ are depleted for genes within the KEGG CAMs pathway module (NES: −1.82; p=0.0078). Bottom: heat map displaying relative expression of individual genes by aging p14 T_M_ (blue: low, red: high). The right hand column summarizes corresponding protein expression by aging splenic or blood-borne D^b^NP_396_^+^ and/or D^b^GP_33_^+^CD8^+^T_M_ retrieved from LCMV-immune B6 mice; colors indicate significant expression changes accrued over time (red: upregulation; black: no change; green: downregulation) and the primary protein expression data in this summary is detailed in panel B as well as ***Figs.5A*, *S2*** and/or refs. [9, 10]. **B.,** temporal regulation of CD11a and CD54 expression by D^b^NP_396_^+^CD8^+^T_E/M_ (triangle symbol: CD44^lo^ CD8^+^T_N_). **C.,** mixed AT/RC Arm and cl13 experiments were performed with rat IgG (control) or aCD11a blocking antibodies; due to the efficacy and prolonged half-life of the KBA antibody, antibodies were only injected on d0 and d2. **D.,** temporal regulation of CXCR3 expression by aging blood-borne D^b^NP_396_^+^CD8^+^T_M_ (dot plots gated on CD8^+^T cells). **E.,** II^o^ CD8^+^T_E_ expansions in different tissues as assessed after LCMV Arm infection and control (hamster Ig) *vs*. αCXCR3 treatment. Note the preferential decrease of aged II^o^ CD8^+^T_E_ expansions in the wake of CD11a or CXCR3 blockade as indicated by black and gray arrows/values (panel C & E) (n≥3 mice/group or time point; AT of 5×10^3^ [panel C top & E] or 8×10^3^ [panel C middle/bottom] young and old D^b^NP_396_^+^CD8^+^T_M_ ach).

LFA-1 biology has been characterized in great detail [37] and the importance of CD11a for the migration of naïve T cells to peripheral LNs is well-established [38], yet the precise role of CD11a in the regulation of pathogen-specific T cell immunity remains incompletely defined. In one of the most detailed report to date, Bose *et al*. found that CD11a-deficiency reduces I^o^ *L. monocytogenes*-(LM-) specific CD8^+^T_E_ responses, skews CD8^+^T_E_ phenotypes in favor of “memory precursors”, preserves major CD8^+^T_E_ functions while stunting *ex vivo* CTL activity, attenuates the subsequent contraction phase, and, remarkably, enhances II^o^ CD8^+^T_E_ expansions by ∼1.8-fold following high-dose LM re-challenge [39]. The latter finding, however surprising, is consistent with our “rebound model” of CD8^+^T_M_ de-differentiation [9, 10] in that any deficits conveyed by CD11a-deficiency are eclipsed by the advanced maturation stage of CD11a^-/-^ CD8^+^T_M_ [39] that is associated with greater recall capacity. Similarly, LFA-1 blockade resulted in a ∼2-fold reduction of I^o^ LCMV-specific CD8^+^T_E_ expansions (ref.[40] and not shown) but its potential impact in the specific context of CD8^+^T_M_ recall responses has not yet been determined. As based on the experience with LFA-1 blockade in transplantation and autoimmunity [41, 42] and considering in particular the lower CD11a and CD54 expression of naïve CD8^+^T_N_ [9], we speculated that CD8^+^T_M_ would be overall more resistant to LFA-1 blockade but that declining CD11a and CD54 levels by aging CD8^+^T_M_ (***Figs.4A/B & S2***) might render them again somewhat more susceptible to this intervention. Using our mixed AT/RC system, LFA-1 blockade in the context of an LCMV cl13 infection indeed promoted a prominent and preferential reduction of aged as compared to young II^o^ CD8^+^T_E_ responses in peripheral blood (4.0-fold *vs*. 2.7-fold) that was less evident in the spleen or after LCMV Arm challenge (***Fig.4C*** and not shown). In fact, blocking LFA-1 in the chronic infection model compromised old CD8^+^T_M_ recall responses to an extent that approaches the decrease observed for concurrent I^o^ CD8^+^T_E_ responses (4.1-fold [p=0.0003] and 2.0-fold [p=0.04] reduction in blood and spleen, respectively). The efficacy of LFA-1 blockade therefore correlates inversely with expression levels of CD11a (and other components of the LFA-1 pathway) on CD8^+^T cells such that the inhibition of proliferative expansion is greater for CD8^+^T_N_ than CD8^+^T_M_, and more substantial for old than young CD8^+^T_M_. We conclude that aged CD8^+^T_M_ populations rely in part on the LFA-1 system to support their improved recall responses in the periphery.

Like the integrins, and often in conjunction, chemokine receptors sensitize T cells to essential spatiotemporal cues required for the effective orchestration of T cell responses [43]. For example, CD8^+^T_E_ and T_M_ subsets can be recruited to reactive LNs by virtue of their CXCR3 expression [44], and several reports have detailed the importance of CXCR3 for the precise positioning of CD8^+^T cells in spleen and LNs, and for the measured rather than accelerated development of I^o^ CD8^+^T_M_ populations [45-48]. However, in regards to the requirement of CD8^+^T_M_-expressed CXCR3 for the regulation of their II^o^ responses, strikingly different conclusions were reached: CXCR3-deficiency either improved II^o^ CD8^+^T_E_ expansions [45], had no effect [46], or compromised II^o^ CD8^+^T_E_ reactivity [48]. The use of different model systems and experimental protocols may have contributed to the divergent outcomes but another factor may be the precise timing of re-challenge experiments since CXCR3 expression by splenic and blood-borne virus-specific CD8^+^T_M_ changes substantially over a period of ∼18 months ([9, 49] and ***Figs.4D & S2***). To circumvent potentially confounding factors associated with the generation of CXCR3^-/-^ CD8^+^T_M_, we used a non-depleting CXCR3 antibody [50] and the results of our mixed AT/RC studies demonstrate that CXCR3 is indeed required for optimal II^o^ CD8^+^T_E_ expansions. Specifically, CXCR3 blockade preferentially weakened the II^o^ response of old as compared to young CD8^+^T_M_, did so in a systemic fashion (i.e. was observed in blood, spleen and LNs), and to an extent that somewhat exceed the impairment of contemporaneous I^o^ CD8^+^T_E_ expansions (2.2-fold [p=0.0005] and 3.5-fold [p=0.0035] decrease in blood and spleen, respectively) (***Fig.4E***). Ready access for CD8^+^T_M_ to local regions of CXCR3 ligand (CXCL9/10) expression [45-48] therefore constitutes an important parameter for the optimal systemic expansion of II^o^ CD8^+^T_E_ populations, and aged CD8^+^T_M_, by virtue of enhanced CXCR3 expression, are poised to effectively harness these interactions.

### CD28- but not CD27-dependent co-stimulation preferentially promotes enhanced II^o^ reactivity of aged CD8^+^T_M_

Recall responses are traditionally regarded as “co-stimulation independent” but more recent work has documented an important role especially for CD28 in the regulation of pathogen-specific II^o^ CD8^+^T_E_ immunity [51]. Although our original analysis of genes differentially expressed by young and old CD8^+^T_M_ included few members of the major co-stimulatory B7 and TNF superfamilies [9], the temporal GSEAs conducted here captured many more subtle alterations, including an upregulation of *Cd28* by aging p14 T_M_ (***Fig.4A*** and not shown). A corresponding age-associated augmentation of CD28 protein expression was confirmed and extended here to blood-borne D^b^NP_396_^+^ and D^b^GP_33_^+^CD8^+^T_M_ populations, and similar experiments corroborated a particularly prominent increase for CD27 (***Figs.5A & S2***), a co-stimulatory receptor that exhibits some of the most pronounced expression differences between young and old CD8^+^T_M_ [9].

**Figure 5.**
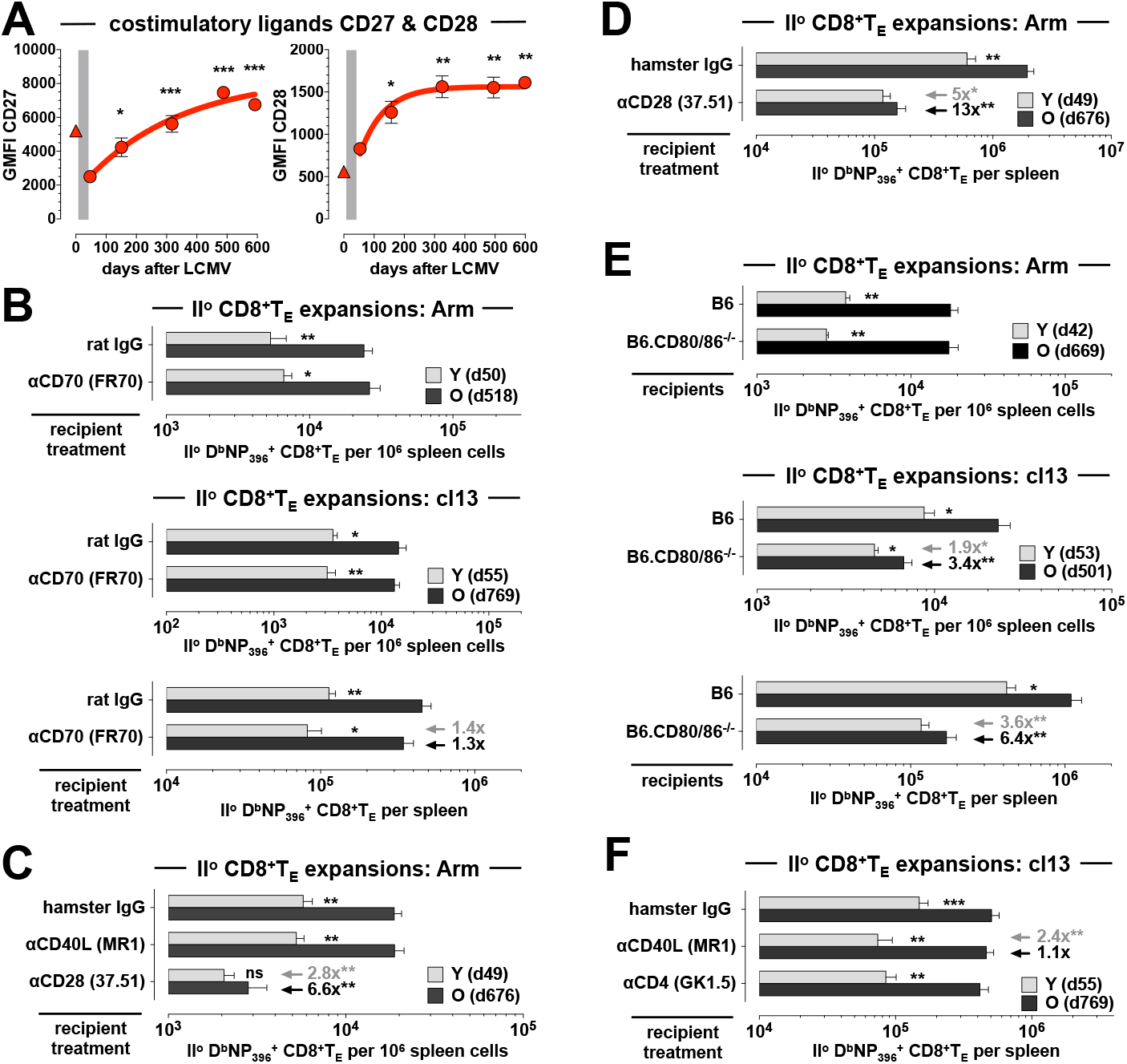
CD28:CD80/86 but not CD27:CD70 or CD40L:CD40 co-stimulatory interactions preferentially promote improved II^o^ reactivity of aged antiviral CD8^+^T_M_. **A.,** temporal regulation of CD27 and CD28 expression by aging D^b^NP_396_^+^CD8^+^T_M_ in peripheral blood (triangle symbol: CD44^lo^CD8^+^T_N_). **B.,** mixed AT/RC experiments were conducted in the LCMV Arm and cl13 systems as indicated using treatment with rat IgG (control) or CD70 blocking antibodies. **C.,** II^o^ CD8^+^T_E_ expansions after mixed AT/RC Arm performed under conditions of CD40L or CD28 blockade. **D.,** same experiment as in C but depicting total splenic II^o^ CD8^+^T_E_ numbers accumulated in the absence *vs*. presence of CD28 blockade. **E.,** mixed AT/RC experiments with CD80/86^-/-^ recipients employing LCMV Arm (top) or cl13 (middle/bottom) infection protocols. The gray and black arrows/values in panels C-D emphasize the extent of reduced II^o^ CD8^+^T_E_ expansions comparing young II^o^ CD8^+^T_E_ populations of control and antibody treated mice (gray), and old II^o^ CD8^+^T_E_ populations of control and antibody treated mice (black); adjacent asterisks indicate statistical significance (n≥3 mice/group or time point; AT of 2×10^3^ [panel B top, C, D & E top], 8×10^3^ [panel B middle/bottom & F] or 10×10^3^ [panel E middle/bottom] young and old D^b^NP_396_^+^CD8^+^T_M_ each).

Despite the general importance of the CD27:CD70 co-stimulatory pathway [52], its contribution to the regulation of LCMV-specific CD8^+^T_E_ immunity appears to be more limited. CD70 blockade or deficiency modestly reduced LCMV-specific I^o^ CD8^+^T_E_ expansions after an acute virus challenge but left the II^o^ response largely intact [53-55]. We made near identical observations in our mixed AT/RC Arm model conducted under conditions of CD70-blockade, *i.e*. we found a small reduction of I^o^ host CD8^+^T_E_ responses (not shown) whereas the overall and differential expansions of young and old II^o^ CD8^+^T_E_ populations were fully preserved (***Fig.5B***). Blocking CD70 in the context of a chronic or high-dose LCMV infection, however, was reported to promote the opposite effect of modestly increasing I^o^ but decreasing II^o^ CD8^+^T_E_ responses [54, 56]. Again, these results were essentially reproduced in our experiments where LCMV cl13-induced I^o^ CD8^+^T_E_ host responses under conditions of CD70 blockade were somewhat elevated (∼1.6-fold, not shown) yet concomitant young and old II^o^ CD8^+^T_E_ expansions were both slightly reduced (***Fig.5B***). Regardless of the relatively small impact exerted by CD70-blockade on the coordination of CD8^+^T_E_ cell immunity, the divergent regulation of I^o^ and II^o^ CD8^+^T_E_ responses in the same microenvironment indicates that CD27:CD70-mediated interactions are not only contingent on pathogen virulence, tropism, persistence and related parameters [52, 53] but also on the differentiation stage of specific CD8^+^T cells themselves. At the same time, the large increase of CD27 expression by aging CD8^+^T_M_ remains unexpectedly inconsequential for the regulation of their II^o^ reactivity.

With regard to the gradual increase of CD28 expression by aging CD8^+^T_M_ (***Figs.4A, 5A & S2***), earlier work by us and others has already implicated the CD28:CD80/86 pathway in the regulation of LCMV-specific II^o^ CD8^+^T_E_ immunity [57, 58] raising the possibility that a more efficient use of these interactions by old CD8^+^T_M_ may boost their recall responses. In confirmation of this prediction, the impairment of II^o^ CD8^+^T_E_ expansions after CD28-blockade in the mixed AT/RC Arm scenario was more pronounced for old as compared to young CD8^+^T_M_ (13-fold *vs*. 5-fold) and resulted in the obliteration of any numerical differences between young and old II^o^ CD8^+^T_E_ populations (***Fig.5C/D***). An accompanying ∼3.5-fold decrease of I^o^ NP_396_-specific host populations (not shown) essentially replicated the phenotype of LCMV-challenged CD28^-/-^ mice [59] and the apparently lesser impact of CD28-blockade on I^o^ CD8^+^T_E_ responses may be due to the lower CD28 expression by CD44^lo^CD8^+^T_N_ (***Fig.5A***). Using an alternative approach to probe the CD28:CD80/86 pathway, we conducted mixed AT/RC experiments with CD80/86^-/-^ recipients. Based on our previous work, we anticipated a critical difference employing LCMV Arm *vs*. cl13 re-challenge protocols: despite the reliance of CD8^+^T_M_ recall responses on CD28, re-challenge with LCMV Arm proved independent of CD80/86 suggesting the existence of another CD28 ligand; in contrast, II^o^ CD8^+^T_E_ expansions were clearly CD80/86-dependent following an LCMV cl13 re-challenge [58]. In agreement with these findings, neither II^o^ nor concurrent I^o^ CD8^+^T_E_ responses elicited in the mixed AT/RC Arm system were affected by CD80/86-deficiency (***Fig.5E***). Yet a LCMV cl13 infection not only reduced CD8^+^T_M_ recall reactivity in general but preferentially comprised the accumulation of aged (6.4-fold) as compared to young (3.6-fold) II^o^ CD8^+^T_E_ (***Fig.5E***). Together, these results support the notion that CD28-mediated co-stimulation contributes specifically to the improved II^o^ reactivity of aged CD8^+^T_M_.

### Role of CD40L and CD4^+^T cells in the differential regulation of young and old II^o^ CD8^+^T_E_ responses

In extension of our investigation into co-stimulatory pathways above, we also evaluated the potential involvement of CD40L:CD40 interactions in the regulation of II^o^ CD8^+^T_E_ immunity, experiments prompted by our observation that aged CD8^+^T_M_ synthesize larger amounts of CD40L upon re-stimulation [9]. Although CD8^+^T cell-produced CD40L appears dispensable for I^o^ CD8^+^T_E_ responses [60], it readily promotes DC activation, B cell proliferation and antibody production (activities usually associated with CD4^+^ “helper” T cells) [61], and may boost II^o^ CD8^+^T_E_ immunity under conditions of limited inflammation [62]. Similarly, our previous work has documented that CD40L blockade administered within the first week of acute LCMV Arm infection does not impinge on I^o^ CD8^+^T_E_ responses but affects subsequent CD8^+^T_M_ development as revealed by impaired II^o^ *in vitro* CTL activity [63]. While these results point towards a more limited and context-dependent role for CD8^+^T cell-produced CD40L, any interpretation of outcomes observed after anti-CD40L treatment has to consider that it targets both CD4^+^ and CD8^+^T cell subsets.

In the mixed AT/RC Arm setting employed here, acute CD40L blockade did not compromise either I^o^ (not shown) or II^o^ CD8^+^T_E_ responses (***Fig.5C***), observations that are also consistent with the finding that provision of additional CD4^+^T cell help did not embellish either I^o^ or II^o^ p14 T_E_ expansions in the wake of an LCMV Arm infection [64]. Yet the situation was reportedly different in the chronic LCMV model: supplementary CD4^+^T cell help increased II^o^ but not I^o^ p14 T_E_ responses, and the effect was abolished by CD40L blockade supporting the conclusion that CD8^+^T_M_ are more reliant than CD8^+^T_N_ on CD40L-mediated CD4^+^T cell help [64]. In the experiments shown in ***Fig.5F***, we quantified CD8^+^T_E_ expansions after mixed AT/RC cl13 under conditions of CD40L blockade. Similar to West *et al*. [64], we found no obvious impact on I^o^ CD8^+^T_E_ responses (not shown) but readily observed a significant reduction of young II^o^ CD8^+^T_E_ populations; nearly identical results were obtained when the experiments were performed with CD4^+^T cell-depleted recipients (***Fig.5F***). Although these results fail to identify a specific contribution for CD8^+^T_M_-expressed CD40L to the regulation of recall responses, they confirm the notion of CD40L:CD40 interactions as an accessory pathway for the optimal elaboration of II^o^ but not I^o^ CD8^+^T_E_ responses. Perhaps most interesting is the fact that aged II^o^ CD8^+^T_E_ reactivity, just like I^o^ CD8^+^T_E_ responses, remained largely unperturbed by either CD40L-blockade or absence of CD4^+^T cell help in the AT/RC cl13 model (***Fig.5F***, and data not shown). This outcome is in fact predicted by the “rebound model” of CD8^+^T_M_ maturation which proposes a progressive harmonization of aging CD8^+^T_M_ properties with those of CD8^+^T_N_ [9, 10], and thus over time a waning importance for CD4^+^T cell help. The model can also explain the seemingly contrasting conclusion that LM-specific CD8^+^T_M_ recall responses become *more* CD4^+^T cell-dependent with age [65]: as opposed to the CD4^+^T cell-independent LCMV Arm system, Marzo *et al*. employed an LM infection protocol where CD4^+^T cell depletion greatly reduced I^o^ CD8^+^T_E_ responses [65]. The complementary observation that CD4^+^T cell depletion in the context of an LM re-challenge also curtailed II^o^ CD8^+^T_E_ expansions, and that this effect became more pronounced with advancing age [65] further indicates that aging CD8^+^T_M_ gradually reestablish a reliance on CD4^+^T cell help akin to that exhibited by CD8^+^T_N_.

### Enforced SAP expression constrains II^o^ CD8^+^T_E_ expansions

The expression patterns of CD2/SLAM family genes and proteins provide yet another example for the converging temporal regulation of CD8^+^T_M_ properties within a defined molecular family: with the exception of stable SLAMF3/Ly9 levels, both mRNA and protein expression of other CD2/SLAM family members were progressively downmodulated in aging CD8^+^T_M_ populations [9]. The functional relevance of these phenotypic changes, however, is difficult to predict since they pertain to both activating and inhibitory receptors, and studies with various blocking/activating antibodies and SLAM receptor-deficient mouse strains have generated at times conflicting results [66]. Even so, since all signaling events transduced by T cell-expressed SLAM receptors operate through the same small adaptor SAP (SLAM-associated protein) [66], a slight decline of *Sh2d1a* message in aging CD8^+^T_M_ was noteworthy [9] in light of earlier work with SAP^-/-^ mice that demonstrated an increase of I^o^ virus-specific CD8^+^T_E_ expansions and associated virus control [66]. The precise cause for this enhancement remains to be determined but a contributing if not essential factor is likely an impairment of activation-induced cell death (AICD) in the SAP^-/-^ mice [67]. In our experiments, however, the decrease of *Sh2d1a* in aging CD8^+^T_M_ was not accompanied by a corresponding reduction of SAP protein expression [9], and eight days after mixed AT/RC Arm, the activation-induced increase of SAP was not significantly different between young and old II^o^ CD8^+^T_E_ (not shown).

Despite these caveats, we chose to explore the additional possibility of differential SAP induction specifically in the earliest phase of the II^o^ response. To this end, we employed the “p14 chimera” model and compared the initial recall response of young and aged p14 T_M_ by CFSE dilution both *in vivo* and *in vitro*. Although we observed similar proliferation patterns for all II^o^ p14 T_E_ populations (***Fig.6A***), more detailed analyses of the *in vitro* studies suggested that aged p14 T_M_ might start to divide a little earlier (*i.e*., exhibiting a ∼1.4-fold higher division indices, not shown) yet the identical proliferation indices of young and old II^o^ p14 T_E_ (***Fig.6A***) are consistent with our earlier conclusion about the comparable antigen-driven proliferation of young and old II^o^ NP_396_-specific CD8^+^T_E_ in the periphery [9]. Importantly though, the better survival of aged II^o^ CD8^+^T_E_ in our *in vivo* model [9] corresponded to higher numbers of old II^o^ p14 T_E_ surviving in the *in vitro* culture system (***Fig.6A***) supporting the general utility of the latter experimental approach. We then proceeded with the quantification of SAP expression as a function of *in vitro* proliferation and found that the early II^o^ effector phase of young but not old p14 T_M_ was accompanied by a significant elevation of SAP levels (***Fig.6A***). Thus, the increased *in vitro* accumulation of aged II^o^ p14 T_E_ correlates with their lower SAP expression which is consistent with the notion of impaired AICD in the absence of SAP [67].

**Figure 6.**
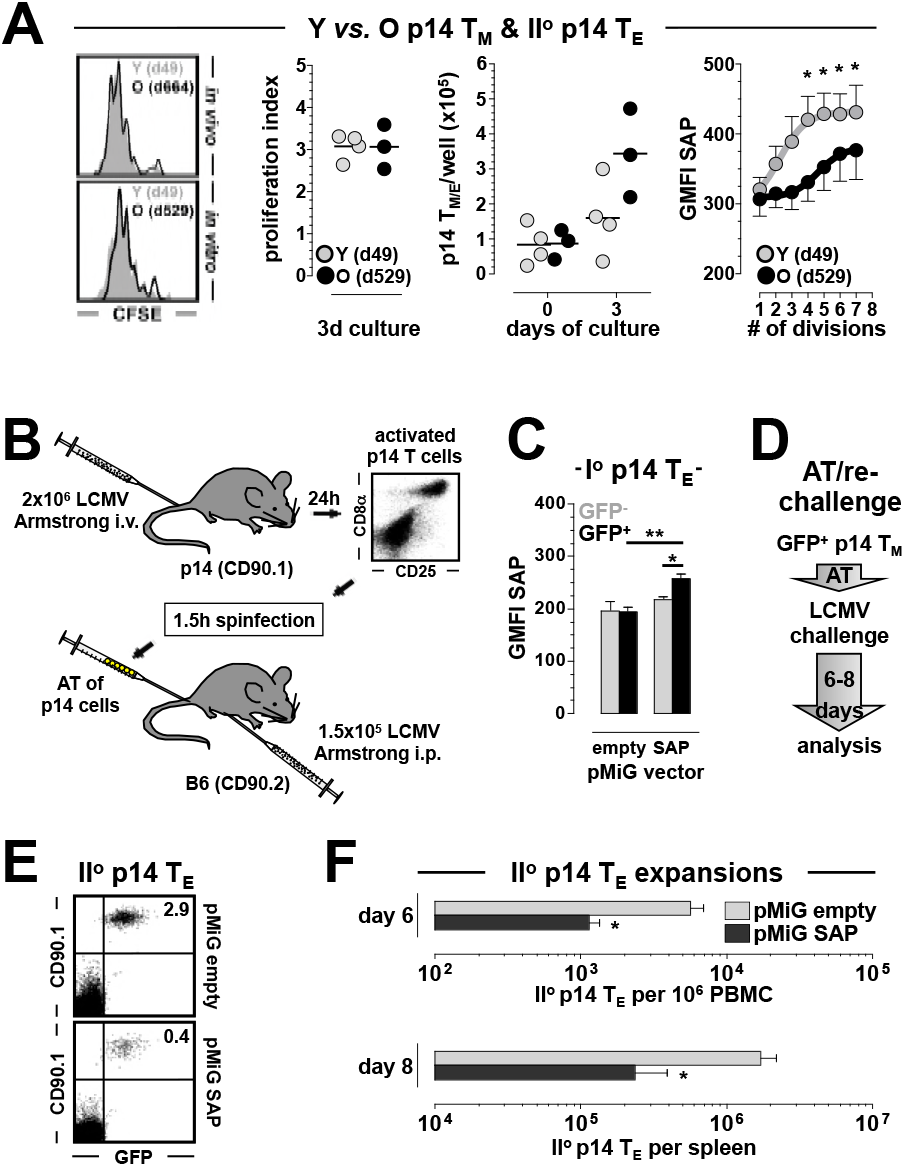
Enforced SAP expression constrains II^o^ reactivity of CD8^+^T_M_. **A.,** proliferation of young and old II^o^ p14 T_E_ as determined by CFSE dilution *in vivo* (64h after AT/RC Arm with 10^6^ p14 T_M_) and *in vitro* (PBMC containing equal numbers of p14 T_M_ cultured for 72h with GP_33-41_ peptide-coated APCs). The adjacent diagrams depict *in vitro* proliferation indices, absolute numbers of p14 T cells at start (d0) and end (d3) of culture, and SAP expression as a function of cell division. **B.,** flow chart for construction of retrogenic p14 chimeras including 1.5h *in vitro* spinfection with pMiG-empty (control) or pMiG-SAP (experimental) retroviruses. **C.,** total SAP content of I^o^ p14 T_E_ (d8) comparing control and experimental p14 chimeras as well as transduced (GFP^+^) and untransduced (GFP^-^) subsets. **D.,** experimental flow chart: GFP^+^ p14 T_M_ (d89) were FACS-purified from control and experimental p14 chimeras and transferred into B6 mice (2×10^3^/recipient) that were then challenged with LCMV Arm and analyzed 6-8 days later. **E.,** II^o^ p14 T_E_ expansions in peripheral blood (d6); dot plots gated on all PBMC, note that GFP expression is restricted to the transferred congenic CD90.1^+^p14 T cells. **F.,** summary of II^o^ p14 T_E_ expansions in blood (d6) and spleen (d8); n≥3 mice/group.

To formally evaluate the hypothesis that the amount of induced SAP expression determines the recall reactivity of II^o^ CD8^+^T_E_ populations, we generated retroviral p14 chimeras that overexpress SAP selectively in subpopulations of p14 T_E/M_ as detailed in ***Fig.6B/C*** and Methods. Following purification of p14 T_M_ transduced with SAP or control retroviruses, AT into naïve B6 hosts and re-challenge with LCMV Arm or cl13 (***Fig.6D***), enforced SAP expression indeed compromised II^o^ p14 T_E_ expansions (***Fig.6E/F***, and data not shown). Collectively, our experiments therefore indicate that the improved antigen-driven II^o^ expansion of aged CD8^+^T_M_ is facilitated by their restrained upregulation of SAP expression. Interestingly, a previous report using T cell hybridomas demonstrated that engagement of a single SLAM receptor, SLAMF4/CD244, could transmit either stimulatory or inhibitory signals depending on the degree of CD244 expression, ligand (CD48) density and, importantly, the level of SAP expression itself [68]. Although those results correlated high SAP expression with activation rather than inhibition [68], an increase of CD244:CD48 interactions could nevertheless convey inhibitory signals even in the presence of elevated SAP expression [69]. Furthermore, in the chronic LCMV model, CD244 was assigned a predominantly inhibitory function as based on enhanced NK cell activity in CD244^-/-^ mice as well as the greater II^o^ reactivity of CD244^-/-^ p14 T_M_ in the AT/RC cl13 system [64, 70], and most recent work specifies that inhibitory functions exerted by the entire *Slam* locus on NK cell responses are solely based on CD244 activity [71]. It should therefore be interesting to assess if the early (though not late) recall response of CD244^-/-^ CD8^+^T_M_ also involves a subdued induction of SAP, and, in more general terms, to determine how precisely CD8^+^T cell-expressed SAP integrates intrinsic signals from SLAMF1-7 receptors [9] *in vivo* to modulate II^o^ CD8^+^T_E_ immunity.

## DISCUSSION

As a widely used preclinical experimental system, the LCMV model has proved extraordinarily helpful in shaping our understanding of protective T cell memory as well as its limitations in the context of persistent viral infections [2, 72, 73]. Here, we used the LCMV model to interrogate over a dozen molecular pathways for their contribution to the embellishment of CD8^+^T_M_ recall responses, and our results are noteworthy for the identification of 1., diverse molecular interactions that specifically promote the greater II^o^ CD8^+^T_E_ expansions of aged CD8^+^T_M_ populations; 2., the distinct outcomes observed after acute *vs*. chronic LCMV re-challenge; and 3., the receptors/ligands that, against expectation, apparently did not participate in the regulation of II^o^ CD8^+^T_E_ immunity.

To explore the possibility that improved recall responses of aged CD8^+^T_M_ emerge as the net result of multiple disparate molecular interactions, we selected a diverse set of pathways comprising cytokine signaling, T cell trafficking, co-stimulation and -inhibition, and effector functionalities as based on the long-term expression kinetics of the respective CD8^+^T_M_-expressed receptors/ligands (the relative robustness of these temporal expression patterns is now supported by an extension of our earlier analyses to blood-borne aging antiviral CD8^+^T_M_ populations, and to subsets with different epitope specificities and TCR affinities/avidities [9]). Our results demonstrate that IL-4-, LFA-1-, CXCR3- and CD28-dependent interactions, a restrained induction of SAP expression, and CD8^+^T_M_-produced FasL and IFNγ not only contribute to efficient II^o^ CD8^+^T_E_ expansions in general, but in particular convey a set of heterogeneous signals that collectively boost the recall reactivity of aged CD8^+^T_M_ populations. While the relative contribution of individual molecular pathways to the regulation of recall responses ranges from the modest to the more pronounced (*e.g*., a 2.8-fold reduction of LCMV Arm-driven aged II^o^ CD8^+^T_E_ expansions under conditions of IFNγ neutralization *vs*. a 13-fold inhibition in the context of CD28 blockade), the overall efficacy of II^o^ CD8^+^T_E_ immunity and immune protection [9] is shaped by the integrated activity of different pathways the individual or combined therapeutic targeting of which may in fact allow for the tailored modulation of specific CD8^+^T_M_ responses.

Furthermore, the above interactions are for the most part of greater importance to the regulation of II^o^ CD8^+^T_E_ immunity in response to a chronic rather than acute viral challenge. Recent work supports the notion that the eventual or at least partial control of chronic viral infections relies on a multiplicity of molecular pathways that are often dispensable for clearance of acute virus infections [74]. Our findings extend this concept to the context of II^o^ CD8^+^T_E_ responses by documenting that CD8^+^T_M_, far from being “co-stimulation independent”, also require the productive engagement of diverse molecular interactions to unfold their full recall potential when confronted with a chronic virus challenge. A further elucidation of these phenomena might very well help to establish an adjusted perspective onto one of the central tenets of T cell memory, namely its presumed imperviousness to the modulation by biochemical pathways commonly referred to as “signal 2 & signal 3”. In fact, the “rebound model” [9], together with the present report, suggests that aging CD8^+^T_M_ become increasingly reliant on the very same “signal 2 & signal 3” interactions that, dependent on the experimental system, also control I^o^ CD8^+^T_E_ differentiation.

Two pathways interrogated in the present study were found to be of preferential importance to the regulation of young rather than old II^o^ CD8^+^T_E_ responses. Here, the greater dependence of young CD8^+^T_M_ recall responses on CD4^+^T cell help and CD40L-mediated interactions in the chronic LCMV system is essentially consistent with the “rebound model”, but the enhanced reliance of young CD8^+^T_M_ on non-CD8^+^T_M_-produced IFNγ, despite reduced CD119/IFNγR1 expression and sensitivity, was unexpected. IFNγ can exert both stimulatory and inhibitory effects on CD8^+^T_E_ populations [29, 75], and the specific balance achieved between these opposing signals may be distinct for young and old CD8^+^T_M_, perhaps as a result of differential IFNγR2 induction [76], but ultimately the reasons for the greater role of host IFNγ in control of young II^o^ CD8^+^T_E_ immunity remain unclear. We also found that several other cytokine signaling and co-stimulatory pathways (IL-6, IL-7, IL-15, TSLP, TGFβ, CD27:CD70) appeared to have at best a minor impact on the regulation of II^o^ CD8^+^T_E_ responses. While these results underscore the obvious fact that promising clues gleaned from a comprehensive set of databases [9] need not necessarily translate into biologically relevant differences within a given model system, they neither rule out potential redundancies not investigated in the present study nor the possibility that these as well as additional pathways may be operative in the context of other experimental and naturally occurring scenarios. Therefore, in as much as the “rebound model” of extended CD8^+^T_M_ maturation applies to pathogen infections in general, the progressive “de-differentiation” of aging CD8^+^T_M_, especially given the “programmed” nature of this process [9], may allow them to brace for more effective recall responses under a greater variety of productive pathogen re-encounters. At the same time, the multitude of diverse molecular pathways involved in shaping improved clinical outcomes also provides an abundance of different targets for potential therapeutic interventions.

## MATERIALS AND METHODS

### Ethics statement

All procedures involving laboratory animals were conducted in accordance with recommendations in the “Guide for the Care and Use of Laboratory Animals of the National Institutes of Health”, the protocols were approved by the Institutional Animal Care and Use Committees (IACUC) of the University of Colorado (permit numbers 70205604[05]1F, 70205607[05]4F and B-70210[05]1E) and Icahn School of Medicine at Mount Sinai (IACUC-2014-0170), and all efforts were made to minimize suffering of animals

### Mice, virus and challenge protocols

C57BL6/J (B6), congenic B6.CD90.1 (B6.PL-*Thy1^a^*/CyJ) and B6.CD45.1 (B6.SJL-*Ptprc^a^ Pepc^b^*/BoyJ), IL-4^-/-^ (B6.129P2-*Il4^tm1Cgn^*/J), IL-6^-/-^ (B6.129S2-*Il6^tm1Kopf^*/J), IFNγ^-/-^ (B6.129S7-*Ifng^tm1Ts^*/J), CD80/86^-/-^(B6.129S4-*Cd80^tm1Shr^Cd86^tm2Shr^*/J) and B6.lpr (B6.MRL-*Fas^lpr^*/J) mice on the B6 background were purchased from The Jackson Laboratory; B6.IL-15^-/-^ (C57BL/6NTac-*IL15^tm1Imx^* N5) mice were acquired from Taconic; and p14 mice harboring TCRtg CD8^+^T cells specific for the dominant D^b^-restricted LCMV-GP_33-41_ determinant were obtained on a B6.CD90.1 background from Dr. M. Oldstone. We only used male mice in this study to avoid potential artifacts that may arise in gender mismatched AT settings. LCMV Armstrong (clone 53b) and clone 13 (cl13) were obtained from Dr. M. Oldstone, and grown and titered as described [77]. For I^o^ challenges, 8-10 week old mice were infected with a single intraperitoneal (i.p.) dose of 2×10^5^ plaque-forming units (pfu) LCMV Arm; for II^o^ challenges, naïve congenic recipients of various CD8^+^T_M_ populations were inoculated with 2×10^5^ pfu LCMV Arm i.p. or 2×10^6^ pfu LCMV cl13 i.v. Infected mice were housed under SPF conditions and monitored for up to ∼2 years. As discussed elsewhere [9, 10], exclusion criteria for aging LCMV-immune mice in this study included gross physical abnormalities (lesions, emaciation and/or weight loss), lymphatic tumors as indicated by enlarged LNs at time of necropsy, and T cell clonal expansions among virus-specific CD8^+^T_M_ populations (D^b^NP_396_^+^, D^b^GP_33_^+^ or D^b^GP_276_^+^); according to these criteria, up to ∼30% of aging mice were excluded from the study.

### Tissue processing, cell purification and adoptive transfers (AT)

Lymphocytes were obtained from blood, spleen and LNs according to standard procedures [78]. Enrichment of splenic T cells was performed with magnetic beads using variations and adaptations of established protocols [9] and reagents purchased from StemCell Technologies, Miltenyi Biotec and Invitrogen/Caltag (***Table S1***). For mixed AT/RC experiments, CD8^+^T_M_ from young and aged LCMV-immune B6 and B6-congenic donors were enriched by combined depletion of B and CD4^+^T cells (or only B cells) followed by 1:1 combination at the level of D^b^NP_396_^+^CD8^+^T_M_, i.v. AT of mixed populations containing 2-10×10^3^ young and old D^b^NP_396_^+^CD8^+^T_M_ each into naïve congenic recipients, and challenge with LCMV (***Figs.1A-D, 2C-E, 3B-D, 4C/E & 5B-F***). For construction of p14 chimeras [9], CD8^+^T cells were enriched from spleens of naïve CD90.1^+^ p14 mice by negative selection, and 5×10^4^ purified p14 cells were transferred i.v. into B6 recipients prior to LCMV infection 2-24h later (***Figs.2B, 3A, 4A, 6A & S1***). *In vivo* proliferation of II^o^ p14T_E_ was assessed by AT of 10^6^ young or old CFSE-labeled p14 T_M_ into B6 recipients and LCMV Arm challenge as detailed in ref [9]; analyses were then conducted with II^o^ p14T_E_ recovered from recipient spleens 64h later (***Fig.6A***).

### Stimulation cultures

Splenic single cell suspensions prepared from young and old LCMV-immune p14 chimeras were cultured for 15min in complete RPMI with graded dosages of recombinant cytokines (murine IL-4, IL-6, IL-10, IFNγ; Peprotech) prior to fixation with PFA buffer, processing and combined CD90.1 and intracellular pSTAT staining (***Figs.2B & 3A***). For *in vitro* proliferation and survival assays, lympholyte-purified PBMC from young and old LCMV-immune p14 chimeras were labeled with CFSE, adjusted to contain the same number of p14 T_M_, and cultured for 72h with T cell-depleted, LCMV-GP_33-41_ peptide-coated B6 spleen cells; numbers of surviving p14 T cells were subsequently calculated using Countess (Invitrogen) or Vi-Cell (Beckmann Coulter) automated cell counters (***Fig.6A***).

### Flow cytometry

All reagents and materials used for analytical flow cytometry are summarized in ***Table S1***, and our basic staining protocols are described and/or referenced in ref.[9]. Detection of phosphorylated STAT proteins (***Figs.2B & 3A***) was performed using a methanol-based cell permeabilization as described [79]. All samples were acquired on FACSCalibur, LSR II (BDBiosciences) or Cyan (Beckman Coulter) flow cytometers and analyzed with DIVA (BDBiosciences) and/or FlowJo (TreeStar) software; visualization of *in vivo* and *in vitro* T cell proliferation by step-wise dilution of CFSE and calculation proliferation and division indices was performed using the FlowJo “proliferation platform” (***Fig.6A***).

### Microarray analyses

Details for microarray analyses of highly purified p14 T_E/M_ populations and time series GSEAs (***Figs.4A & S1***) are found in refs.[9, 10].

### In vivo *antibody treatment*

For *in vivo* blockade of cytokine signaling, T cell trafficking or co-stimulation (***Figs.1B/C, 2E, 3B/C, 4C/E & 5B-D/F***), naïve B6 or B6 congenic recipients were injected i.p. with antibodies ∼2h before AT of mixed CD8^+^T_M_ populations and subsequent LCMV infection as well as on d2 and d4 after challenge (αIL-7 [M25] & αIL-7Ra [A7R34]: 3×500µg each; aTGFp_12_,3 [1D11.16.8]: 3×1000µg; aIFNy [XMG1.2]: 3×1000µg; αCXCR3 [CXCR3-173]: 3×100µg; αCD70: [FR70]: 3×250µg; αCD154/CD40L [MR1]: 3×250µg; αCD28 [37.51]: 3×100µg; αCD62L [MEL-14]: 3×150µg); αCD11a/LFA-1 [KBA]: 2×200µg on d0 and d2 only; corresponding control antibodies: dosages commensurate to experimental antibodies); CD4^+^T cell depletion was achieved by i.p. injection of 200µg GK1.5 antibody on days −1 and +1 in relation to AT/RC [80]. Further details about all *in vivo* antibodies are provided in ***Table S1***.

### Retroviral transductions, chimera construction and transduced p14 T_M_ purification

Murine *Sh2d1a* (SAP) cDNA was purchased from Open Biosystems (clone ID 1400188) and sub-cloned into a murine stem cell virus- (MSCV-) based retroviral pMiG vector that contains GFP as a reporter (gift from P. Marrack). To generate retroviruses, pMiG-empty or pMiG-SAP plasmids were co-transfected with PsiEco helper plasmid into Phoenix 293T cells using Fugene 6 (Roche) according to standard procedures [79]. After 48h, retroviral supernatants were harvested and spin-transductions of *in vivo* activated p14 splenocytes (naïve p14 mice infected with 2×10^6^ pfu LCMV Arm i.v. 24h earlier) were performed for 90min at 32°C in the presence of 8µg/mL polybrene, 10mM HEPES and 10µg/mL recombinant hIL-2. Transduced p14 splenocytes were transferred “blind” into naïve B6 mice that were subsequently infected with 2×10^5^ pfu LCMV Arm i.p. (***Fig.6B***), and effective transduction levels were verified in blood-borne p14 T_E_ 8 days later (***Fig.6C***). For subsequent AT/RC experiments, transduced p14 T_M_ (CD4-B220-CD90.1^+^GFP^+^) were purified from spleens using a Coulter Moflo XDP cell sorter.

### Statistical analyses

Data handling, analysis and graphic representation was performed using Prism 6.0c (GraphPad Software). All data summarized in bar and line diagrams are expressed as mean ±1 standard error (SEM), and asterisks indicate statistical differences calculated by Student’s t-test, or one-way ANOVA with Dunnett’s multiple comparisons test, and adopt the following convention: *: p<0.05, **: p<0.01 and ***: p<0.001.

## ACKNOWLEDGEMENTS

We wish to thank Drs. R. Gill, P. Marrack and A. Veillette for the generous gift of several unique antibodies (***Table S1***), Drs. Z. Yi and W. Zhang for conducting the GSEAs, Dr. E. Clambey for assistance with the collection of blood and tissue samples, and the NIH Tetramer Core Facility for provision of biotinylated MHC:peptide monomers. This work was supported by NIH AG026518 and AI093637, JDRF CDA 2-2007-240, and DERC P30-DK057516 (DH), and NIH T32 training grants AI07405, AI052066 and DK007792 (BD); the funders had no role in study design, data collection and analysis, decision to publish, or preparation of the manuscript.

---

**Figure S1.**
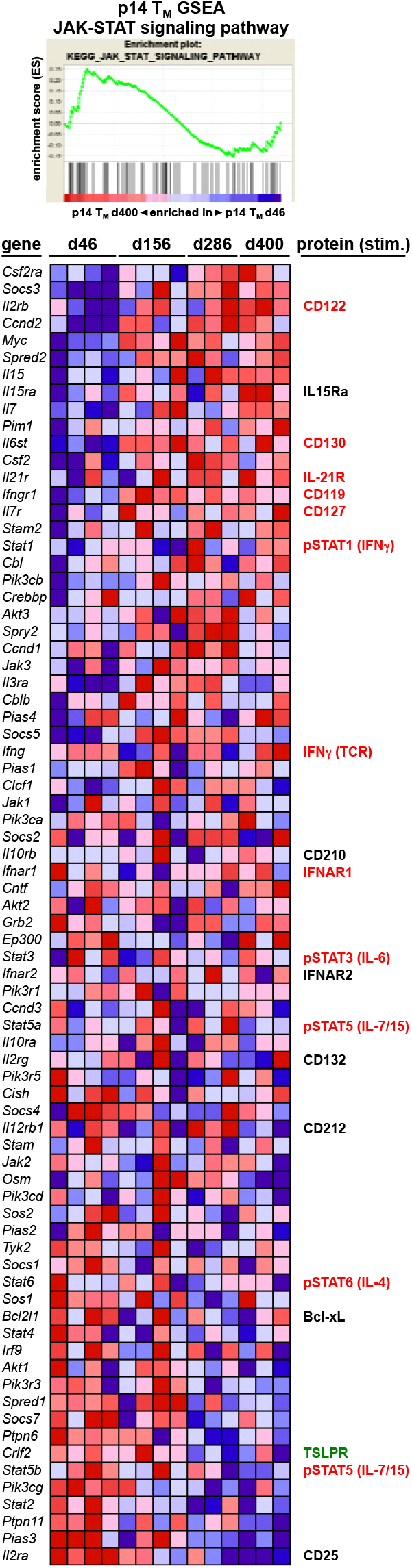
Gene set enrichment analysis (GSEA) of aging p14 T_M_: the JAK-STAT signaling pathway. Time series GSEA were conducted with data sets obtained for aging p14 T_M_ (d46, d156, d286 and d400) purified from LCMV-challenged p14 chimeras and processed directly ex vivo for microarray hybridization as detailed in refs. [9, 10]. Top: aged p14 T_M_ are enriched for genes within the KEGG JAK-STAT signaling pathway module (normalized enrichment score: 1.05). Bottom: heat map displaying relative expression of individual genes by aging p14 T_M_ (blue: low, red: high). The right hand column summarizes corresponding protein expression patterns conducted with aging D^b^NP_396_^+^ and/or D^b^GP_33_^+^ CD8^+^T_M_ retrieved from spleen or blood of LCMV-immune B6 mice; colors identify significant expression changes accrued over time (red: upregulation; black: no change; green: downregulation); where indicated in parenthesis, CD8^+^T_M_ were stimulated in vitro prior to analysis (IFNγ: 5h TCR stimulation with peptide; phosphorylated STAT proteins: 15min stimulation with indicated cytokines). The primary protein expression data in this summary are shown in ***Figs.2A/B, 3A, S2*** and/or refs. [9, 10].

**Figure S2.**
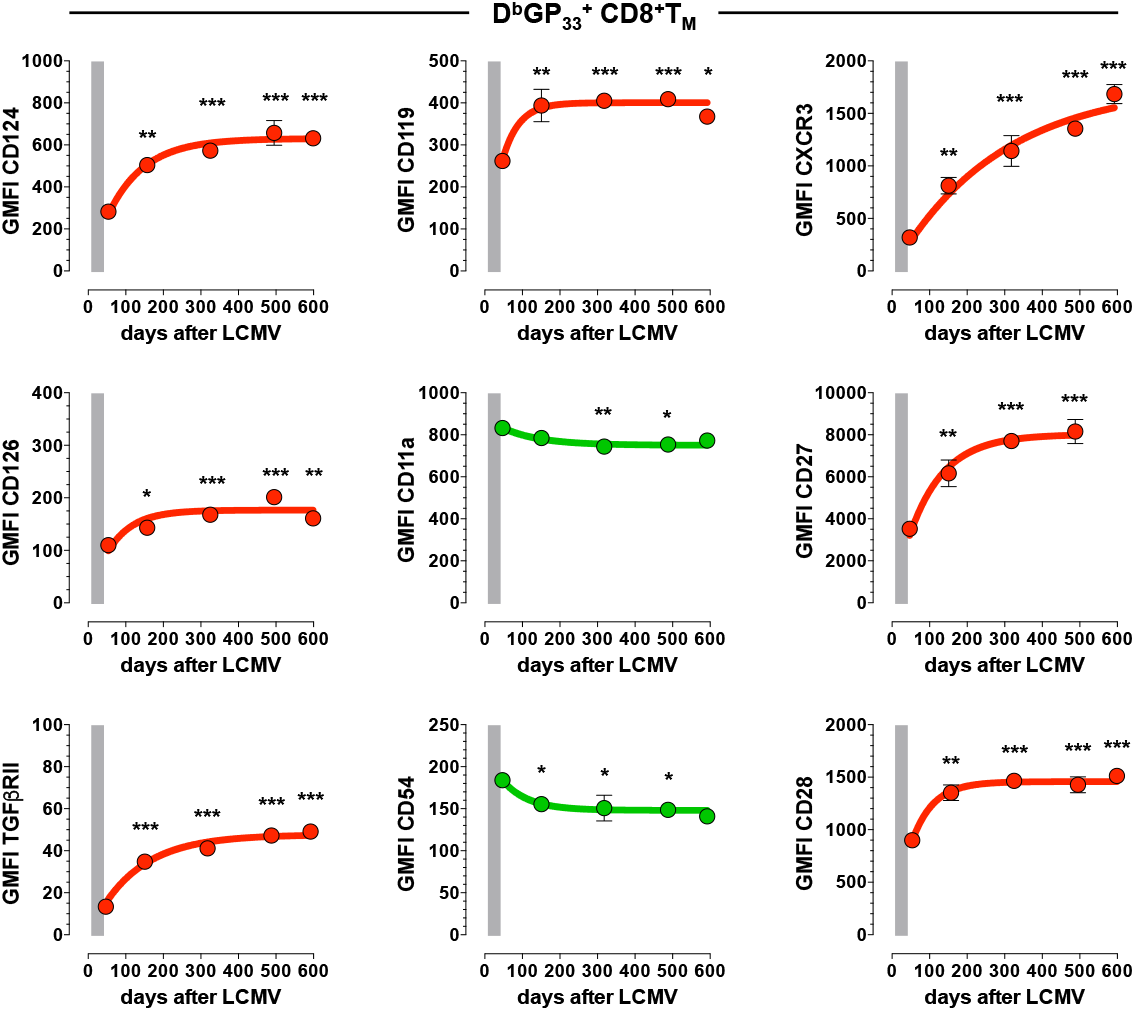
Temporal regulation of selected CD8^+^T_M_-expressed cell surface receptors/ligands. PBMC obtained from cohorts of aging LCMV-immune mice were contemporaneously stained to quantify expression levels of indicated receptors/ligands by D^b^GP_33_^+^ CD8^+^T_M_ (GMFI: gometric mean of fluoresecence intensity; n≥3 mice per time point; statistical differences between young and older CD8^+^T_M_ were calculated using oneway ANOVA with Dunnett’s multiple comparisons test).

## ABBREVIATIONS

AICD: activation-induced cell death
AT: adoptive transfer
AT/RC: adoptive transfer/re-challenge
AT/RC Arm: adoptive transfer/re-challenge with LCMV Armstrong
AT/RC cl13: adoptive transfer/re-challenge with LCMV cl13
BMP: bone morphogenetic protein
CTL: cytotoxic T lymphocyte
GFP: green-fluorescent protein
GP, NP: glycoprotein, nucleoprotein
GSEA: gene set enrichment analysis
I^o^, II^o^, III^o^: primary, secondary, tertiary
KEGG: Kyoto Encyclopedia of Genes and Genomes
LCMV: lymphocytic choriomeningitis virus
LCMV Arm: LCMV Armstrong (clone 53b)
LCMV cl13: LCMV clone 13
LM: *Listeria monocytogenes*
LN: lymph node
O, Y: old, young
p14 (TCRtg): TCRtg CD8^+^T cells specific for the LCMV-GP_33-41_ determinant
PBMC: peripheral blood mononuclear cells
pfu: plaque forming units
SAP: SLAM-associated protein
SLAMF: signaling lymphocyte activation molecule family
STAT, pSTAT: signal transducer and activator of transcription, phosphorylated STAT
T cell subsets: 
T_E_: effector T cells
T_EM_: effector memory T cells (CD62L^lo^/CCR7^-^)
T_CM_: central memory T cells (CD62L^hi^/CCR7^+^)
T_M_: memory T cells
T_MP_: memory-phenotype T cells (CD44^hi^)
T_N_: naïve T cells (CD44^lo^)
T_REG_: regulatory T cells
TCRtg: T cell receptor transgenic
TSLP: Thymic stromal lymphopoietin

